# The planktonic protist interactome: where do we stand after a century of research?

**DOI:** 10.1101/587352

**Authors:** Marit F. Markussen Bjorbækmo, Andreas Evenstad, Line Lieblein Røsæg, Anders K. Krabberød, Ramiro Logares

## Abstract

Microbial interactions are crucial for Earth ecosystem function, yet our knowledge about them is limited and has so far mainly existed as scattered records. Here, we have surveyed the literature involving planktonic protist interactions and gathered the information in a manually curated *Protist Interaction DAtabase* (PIDA). In total, we have registered ~2,500 ecological interactions from ~500 publications, spanning the last 150 years. All major protistan lineages were involved in interactions as hosts, symbionts, parasites, predators and/or prey. Symbiosis was the most common interaction (43% of all records), followed by predation (39%) and parasitism (18%). Using bipartite networks, we found that protistan predators seem to be “multivorous”, while parasite-host and symbiont-host interactions appear to have moderate degrees of specialization. The SAR supergroup (i.e. Stramenopiles, Alveolata and Rhizaria) heavily dominated PIDA, and comparisons against a global-ocean molecular survey (*TARA Oceans*) indicated that several SAR lineages, which are abundant and diverse in the marine realm, were underrepresented among the compiled interactions. All in all, despite historical biases, our work not only unveils large-scale eco-evolutionary trends in the protist interactome, but it also constitutes an expandable resource to investigate protist interactions and to test hypotheses deriving from omics tools.

## Introduction

Aquatic microbes, including unicellular eukaryotes (protists) and prokaryotes, are essential for the functioning of the biosphere^1–4^. Microbes exist in diverse ecological communities where they interact with each other as well as with larger multicellular organisms and viruses. Interaction between microbial species has played important roles in evolution and speciation. One of the best examples is that the origin of eukaryotes is grounded in the interaction-events of endosymbiosis; giving rise to mitochondria, chloroplasts and other metabolic capacities in the eukaryotic cell^5–8^. Microbial interactions guarantee ecosystem function, holding crucial roles in for instance carbon channelling in photosymbiosis, control of microalgae blooms by parasites, and phytoplankton-associated bacteria influencing the growth and health of their host. Despite their importance, our understanding of microbial interactions in the ocean and other aquatic systems is rudimentary, and the majority of them are still unknown^4,9–11^. The earliest surveys of interactions between aquatic microbes date back to the 19^th^ century. In 1851, while on board H.M.S *Rattlesnake* in the Pacific Ocean, Thomas Huxley discovered small yellow-green cells inside the conspicuous planktonic radiolarians which he thought were organelles^12^. Later on, Karl Brandt established that the yellowish cells were symbiotic alga and named them *Zooxanthella nutricola^13^*. Since these early studies, hundreds of others have reported microbial interactions by using classic tools, mainly microscopy, but this knowledge has not yet been gathered in one accessible database. More recently, methods such as environmental High-Throughput Sequencing (HTS)^14^ of DNA or RNA started to transform our understanding of microbial diversity, evolution, and ecological interactions. HTS-based studies have generated hypotheses on microbial interactions based on correlations^15–18^, by exploring the abundance of genetic markers from microbes. These hypotheses need to be tested with other type of data, such as known associations from the literature^19^.

Here, our main objectives were to compile the knowledge on aquatic protist interactions from the literature and make it available to the scientific community. We also report the main patterns found in this survey. We examined the available scientific literature spanning the last ~150 years, and recorded ~2,500 ecological interactions from ~500 publications going back to the late 1800’s^20^ (Supp. Fig.1). Based on this, we generated a manually curated and publicly available *Protist Interaction DAtabase* (PIDA; DOI: 10.5281/zenodo.1195514). PIDA entries have been grouped into three types of pairwise ecological interactions: *parasitism, predation and symbiosis*. Parasitism is an antagonistic relationship between organisms which is beneficial to one but harmful to the other, predation refers for the most part to the engulfment of smaller cells through phagocytosis, and symbiosis refers to interactions beneficial for both partners (mutualism) or beneficial for one and neutral for the other (commensalism).

The taxonomic classification in PIDA includes genus and species level, in addition to three levels that were chosen pragmatically to make the database more user-friendly and portable. The highest level distinguishes between eukaryotic and prokaryotic taxa. The second level places each taxon within supergroups or other high taxonomic levels (e.g. Rhizaria, Alveolata) following the scheme of Adl et al^21,22^, and will be referred to as ‘supergroup level’ in this study. The third level places each taxon in larger groups below the supergroup taxonomic levels (e.g. Ciliophora, Dinoflagellata and Acantharia), and will here be referred to as ‘phylum level’.

## Results

### Aquatic microbial interactions

The literature in PIDA was dominated by studies based on direct observation of interactions such as light microscopy. In total, 82% of the entries were based on microscopy, and only 38% of those were combined with molecular methods. The most commonly studied interaction in the literature was symbiosis, representing 43% of all entries, followed by predation (39%) and parasitism (18%).

The SAR supergroup (Alveolata, Stramenopiles and Rhizaria) dominated with ~92% of the total entries (Figs. 1 & 2). Of all host and predator records, ~90% belonged to the SAR supergroup (Alveolata 51%, Stramenopiles 12% and Rhizaria 27%; Fig. 2). The SAR supergroup was less dominant as symbiont/ parasite/ prey, but still represented the largest group, with 50% of all entries (Alveolata 33%, Stramenopiles 16% and Rhizaria 1%; Fig. 2). Within the SAR supergroup the well-known and species rich Diatomea (Stramenopiles), Dinoflagellata and Ciliophora (both Alveolata) dominated (Fig. 5).

**Fig. 1.**
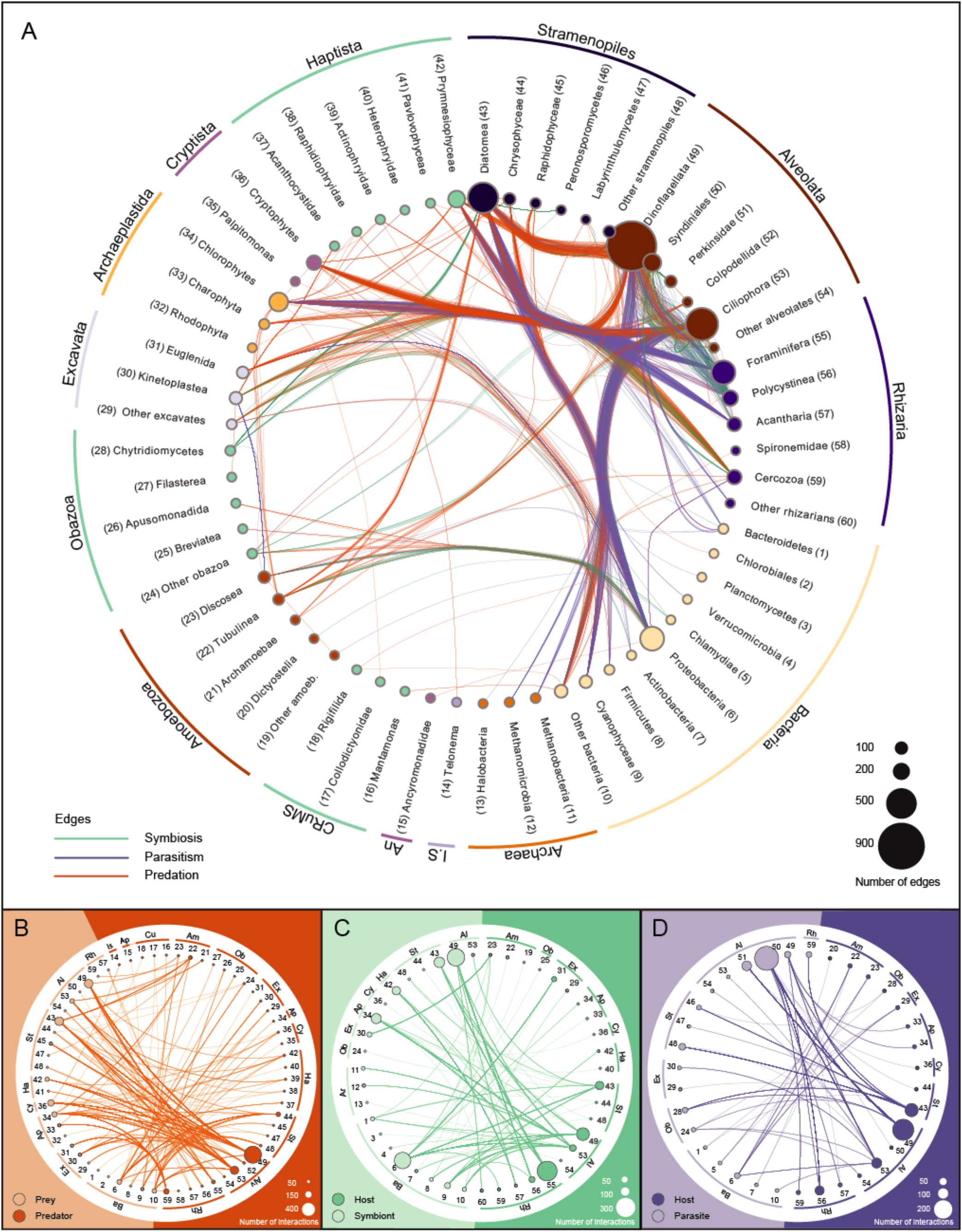
Overview of the interactions included in PIDA. Nomenclature and taxonomic order of Eukaryota is based on Adl et al. 2018^22^. Nomenclature and taxonomic order of Bacteria is based on Schultz et al. 2017^72^. The nodes are grouped (outer circle) according to eukaryotic supergroups (or *Incertae sedis*), Bacteria and Archaea. **Panel A**: Network based on the 2,422 entries in the PIDA database. Nodes represent eukaryotic and prokaryotic taxa and are coloured accordingly. Node size indicates the number of edges/links that are connected to that node. Each node/taxon is assigned a number, which corresponds with the numbers for taxa in B-D. Edges represent interactions between two taxa and are coloured according to ecological interaction type: *predation* (orange), *symbiosis* (green) and *parasitism* (purple). The network is undirected, meaning that a node can contain both parasites/symbionts/prey and hosts/predators. To avoid cluttering of the figure, ‘Self-loops’, which represent cases where both interacting organisms belong to the same taxon (e.g. a dinoflagellate eating another dinoflagellate) are not shown as edges/links in this figure, but are considered in the size of nodes, which represent total number of edges for each node including ‘self-loops’. The outermost circle groups taxa in the different eukaryotic ‘‘supergroups” or the prokaryotic domains Bacteria and Archaea. Ancryomonadidae is abbreviated An. Taxa that are not placed into any of the supergroups are grouped together as *Incertae sedis* (abbreviated *I.S.* in the figure). In **Panels B, C, D** the following abbreviations for supergroups are used: Ar = Archaea, Ba = Bacteria, Rh = Rhizaria, Al = Alveolata, St = Stramenopiles, Ha = Haptista, Cy = Cryptista, Ap = Archaeplastida, Ex = Excavata, Ob = Obazoa, Am = Amoebozoa, Cu = CRuMS, An = Ancryomonadidae, Is = *Incertae sedis*. **Panel B: Predator-prey interactions in PIDA**. The taxa node numbers correspond to numbers in Panel A. Abbreviations for supergroups are described above. Background and nodes are coloured according to functional role in the interaction: Prey are coloured light orange (left part of figure), while predators are depicted in dark orange (right part of figure). The size of each node represents the number of edges (entries in the database) connected to that node. **Panel C: Symbiont - host interactions included in PIDA**. The node numbers correspond to node numbers in Panel A. Abbreviations for supergroups are described above. Symbionts are to the left, coloured light green, and their hosts are to the right in dark green. The size of each node represents the number of edges (entries in the database) connected to that node. **Panel D: Parasite - host interactions included in PIDA**. The node numbers correspond to node numbers in Panel A. Abbreviations for supergroups are described above. Parasite taxa are depicted in light purple (left), hosts in dark purple (right).

**Fig. 2.**
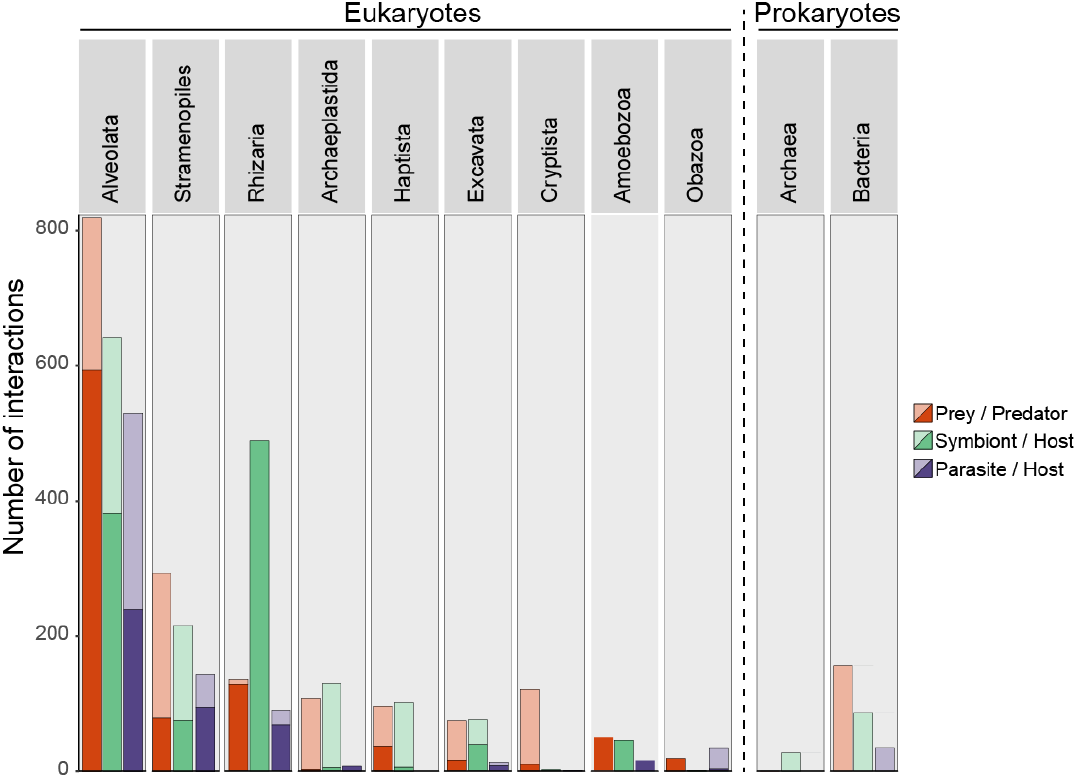
Interactions in PIDA. Number of interactions registered in the PIDA database for the different taxonomic groups at ‘supergroup level’ (corresponding to the second taxonomic level in PIDA). Red bars show predation, green represent symbiosis and purple parasitism. Solid colours represent host/predator and transparent colours represent symbiont/parasite/prey. Because CRuMS, Ancyromonadidae and *Incertae sedis* comprised very few entries (5, 1 and 2 predator entries respectively), they are not included in this figure.

The majority of interactions (82%) were from marine or brackish waters, while studies from freshwater systems accounted for a smaller fraction of the interactions (18%). This is not surprising given the larger number of studies from the marine phototrophic zone compared to other environments.

### Predator – prey interactions

Protist predation can be divided into different categories depending on the type of prey involved: herbivory (grazing on autotrophic eukaryotic algae), bacterivory (feeding on autotrophic and/or heterotrophic bacteria) or predation on other heterotrophic protists. Herbivory was the most common form of predation in our survey with all the major eukaryotic lineages represented among the predators. Entries of herbivore dinoflagellates and ciliates (both Alveolata) dominated (Fig. 1A-B). Bacterivory accounted for 16% of the predator – prey interactions and was also documented in most eukaryotic groups (Fig. 1B), with an expected predominance of small heterotrophic flagellates.

### Symbiont – host interactions

Symbiont interactions constituted the majority of entries in PIDA. Symbiotic *protist-protist* interactions made up 63% of the symbiont entries and the majority of these interactions represented photosymbiosis (86%, while the remaining 14% were classified as “unknown symbiotic relationship”). Dinoflagellates, diatoms, chlorophyceans, trebouxiophyceans, and prymnesiophytes accounted for most of the recorded photosymbionts, living in symbiosis with rhizarian, ciliate and dinoflagellate hosts (Fig. 1A, C). *Bacteria-protist* interactions represented 34% of the total number of symbiont entries in PIDA, and was dominated by bacterial entries belonging to Proteobacteria, Cyanophyceae and Bacteroidetes that interacted with the majority of protist supergroups (except Cryptista, CRuMS, Ancyromonadidae and Telonema; Fig. 1A, C). The bacteria-protist interactions were involved in many different types of symbiotic relationships, from photosymbiosis (4%) to nitrogen fixation (14%) and vitamin exchange (11%). The majority of the bacteria-protist relationships were, however, classified as “unknown symbiotic relationship” (69%). Symbiotic *archaea-protist* interactions represented 3% of symbiont entries in PIDA, and the majority of these were methanogenic symbiont interactions between archaeal Metanomicrobia and anaerobic Ciliophora (Fig. 1A, C).

### Parasite – host interactions

Parasites in PIDA were dominated by a few taxonomic groups that all belonged to Alveolata, such as Syndiniales (~50% *Amoebophrya*), Perkinsidae (~98% *Parvilucifera*) and Dinoflagellata. Together they accounted for 2/3 of the parasite entries (Fig. 1 A, D). These alveolate parasites mainly infected other alveolates such as dinoflagellates and ciliates, but rhizarian and diatom hosts were also recorded (Fig. 1D). Parasites belonging to different stramenopiles lineages such as Peronosporomycetes (oomycetes), Labyrinthulomycetes and Pirsonia were mainly described from diatom hosts (Fig. 1D). Among rhizarian parasites there were just a few cercozoans and phytomyxids (5% of the parasites) that were registered to parasitize other protists (diatoms and one species of Perenosporomycetes, both Stramenopiles). Parasitic fungi from Chytridiomycetes, Microsporidia and Sordariomycetes (the last two included in “other Obazoa” in Fig. 1D) were also represented by relatively few entries (only 7% of the parasite records). Yet, the records of parasitic fungi demonstrated that they infect a relatively broad range of protists, such as dinoflagellates, apicomplexans, ciliates and diatoms. Bacterial parasites of protists accounted for 8% of the parasite entries and were registered mainly from amoebozoan, excavate and ciliate hosts (Fig. 1D).

### Bipartite interaction networks

Since PIDA consists of pairwise interactions between aquatic microbes where the roles of the participants are known we can represent the interactions as bipartite networks. Bipartite networks provide a systematic way of representing data that consist of two distinct guilds, such as plant-pollinator, parasite-host or predator-prey. These networks are composed of nodes (representing species or genera) connected by links (edges) representing the interactions between nodes. The degree of a node (species) is the sum of links connecting the particular node to the nodes from the other guild. Consequently, a higher degree value indicates a higher level of generalism^23^. For example, a parasite that has gone through multiple host-shifts and has the capacity to parasitize different hosts would display a higher degree than a parasite specialized to interact with only one host organism. We have constructed binary (presence/absence) bipartite networks for predator-prey, symbiont-host and parasite-host interactions to characterize and compare the three types separately (Supplementary Figs. 2–4). We calculated specialization indices to analyse variation in specialization within the bipartite networks and to examine if the three interaction types differed in terms of specialization (Fig. 3 A-D; Table 1).

**Table 1:** Degree and specialization index (d’), for the five species with highest degree (highest number of links/edges) in the bipartite networks for each interaction type. #Taxa displays the number of species registered in PIDA for the different interaction types. The taxonomy of the five species with highest degree is shown at species, phylum and supergroup level. Degree shows the number of edges for the top five taxa. Specialization index *d*’ (Kullback-Leibler distance)^68^ ranges from 0 for the most generalized to 1 for the most specialized. Abbreviations used: Type = Interaction type, symb = symbionts, par = parasites.

| Type | #Taxa | Species | Phylum (Supergroup) | Degree | $d'$ |
| --- | --- | --- | --- | --- | --- |
| Predators | 337 | *Leptophrys vorax* | Cercozoa (Rhizaria) | 34 | 0.8 |
|  |  | *Oblea rotunda* | Dinoflagellata (Alveolata) | 27 | 0.6 |
|  |  | *Karlodinium armiger* | Dinoflagellata (Alveolata) | 26 | 0.6 |
|  |  | *Diplopsalis lenticula* | Dinoflagellata (Alveolata) | 15 | 0.4 |
|  |  | *Diplopsalopsis bomba* | Dinoflagellata (Alveolata) | 15 | 0.4 |
| Hosts of symb | 309 | *Amphistegina lobifera* | Foraminifera (Rhizaria) | 17 | 0.7 |
|  |  | *Amphistegina lessonii* | Foraminifera (Rhizaria) | 15 | 0.6 |
|  |  | *Borelis schlumbergi* | Foraminifera (Rhizaria) | 12 | 0.6 |
|  |  | *Heterostegina depressa* | Foraminifera (Rhizaria) | 12 | 0.5 |
|  |  | *Paramecium bursaria* | Ciliophora (Alveolata) | 9 | 0.8 |
| Hosts of par | 262 | *Alexandrium minutum* | Dinoflagellata (Alveolata) | 5 | 0.3 |
|  |  | *Collozoum* NA | Polycystinea (Rhizaria) | 5 | 0.8 |
|  |  | *Guinardia delicatula* | Diatomea (Stramenopiles) | 5 | 0.6 |
|  |  | *Alexandrium catenella* | Dinoflagellata (Alveolata) | 4 | 0.1 |
|  |  | *Coscinodiscus granii* | Diatomea (Stramenopiles) | 4 | 0.5 |
| Prey | 342 | *Isochrysis galbana* | Prymnesiophyceae (Haptista) | 31 | 0.3 |
|  |  | *Rhodomonas salina* | Cryptophyceae (Cryptista) | 27 | 0.4 |
|  |  | *Skeletonema costatum* | Diatomea (Stramenopiles) | 27 | 0.3 |
|  |  | *Heterocapsa triquetra* | Dinoflagellata (Alveolata) | 20 | 0.3 |
|  |  | *Heterosigma akashiwo* | Raphidiophyceae (Stramenopiles) | 19 | 0.2 |
| Symbionts | 167 | *Phaeocystis* NA | Prymnesiophyceae (Haptista) | 29 | 1.0 |
|  |  | cyanophyte NA | Cyanophyceae (Bacteria) | 25 | 0.9 |
|  |  | gammaproteobacteria NA | Proteobacteria (Bacteria) | 24 | 0.7 |
|  |  | alphaproteobacter NA | Proteobacteria (Bacteria) | 23 | 0.6 |
|  |  | diatom NA | Diatomea (Stramenopiles) | 23 | 1.0 |
| Parasites | 130 | *Amoebophrya* NA | Syndiniales (Alveolata) | 61 | 0.8 |
|  |  | *Parvilucifera infectans* | Perkinsidae (Alveolata) | 29 | 0.5 |
|  |  | *Euduboscquella* NA | Syndiniales (Alveolata) | 18 | 1.0 |
|  |  | *Cryothecomonas longipes* | Cercozoa (Rhizaria) | 14 | 0.6 |
|  |  | *Pirsonia formosa* | I .s . Stramenopiles | 14 | 0.6 |

**Fig. 3.**
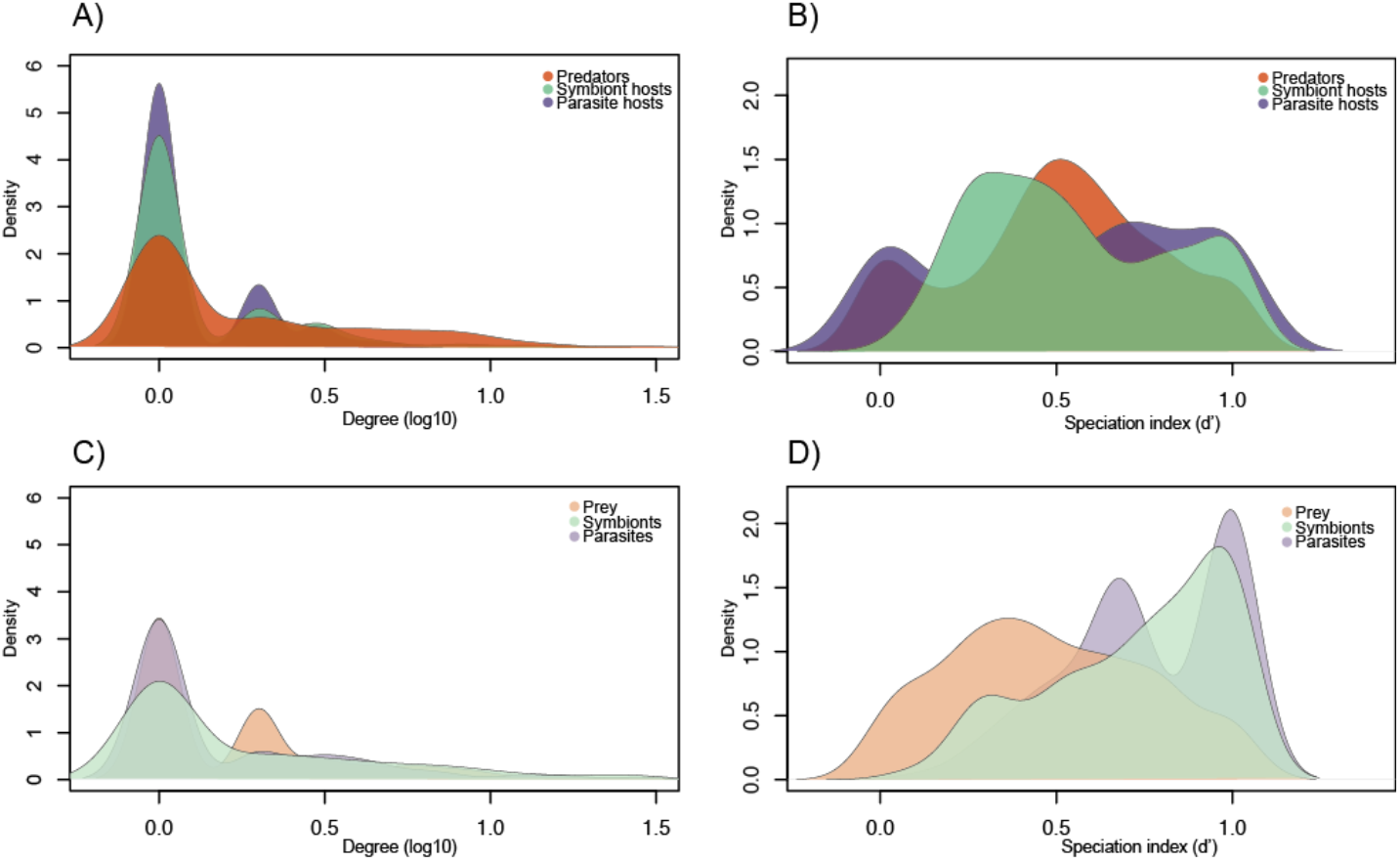
Degree and Specialization indices for the bipartite networks in PIDA. **A**) Degree (number of links/edges/interactions) for predators, host of parasites and host of symbionts in the bipartite networks. **B**) Density plot of the Specialization index *d*’ (Kullback-Leibler distance)^70^, for predators, host of parasites and host of symbionts. The specialization index *d*’ ranges from 0 for the most generalized to 1 for the most specialized. **C**) Degree (number of links/edges/interactions) for prey, parasites and symbionts. **D**) Density plot of the Specialization index *d*’ for prey, parasites and symbionts.

The *predator-prey* bipartite interaction networks had 342 prey species and 337 predator species (Table 1). Although the number of prey and predators in the network were almost equal there are multiple shared interactions. That is, several predators can feed on the same prey (i.e. the prey has a high degree) and conversely, predator species with a high degree indicates taxa that are generalist preying on multiple prey organisms (Fig 3 A and C). The five predators with highest degree included four dinoflagellates and one cercozoan (Table 1). The prey organisms with highest degree values belonged to Haptista, Cryptista, Stramenopiles and Alveolata (Table 1). The specialisation index (*d*’) was uniformly distributed from 0 (generalist) to 1 (specialist) indicating that predation was not pushing predator or prey to specialisation (Fig 3 B and D).

The *symbiont-host* interaction networks consisted of 167 symbiont and 309 host species (Table 1). The majority of both hosts and symbionts had a low degree (Fig. 3 A and C). The distribution of the specialisation index (*d*’) for hosts indicates that PIDA includes both specialists interacting with only one or a few symbionts as well as those that interact with multiple symbionts (Fig. 3B). The five host taxa that had the highest number of associated symbionts (i.e. highest degree) were four foraminiferans (Rhizaria) and one ciliate (Alveolata; Table 1). Very few hosts were “perfect generalists” (i.e. with *d*’ close to 0, Fig. 3B). The symbionts in PIDA had high *d*’ values in general, which indicates that they are more specialised (*d*’ 0.75-1; Fig. 3D). The degree also shows that most symbionts have few links to different host species (Fig. 3C).

The network for *parasitism* had 130 parasites and 262 hosts (Table 1). Hosts were dominated by taxa with low degree (i.e. few parasites per host), which indicated that they are infected by a relatively low number of parasites per host (Fig. 3A). The *d*’ values showed, however, that there was an equal distribution of host taxa ranging from “perfect generalists” (*d*’ value of 0) to “perfect specialists” (*d*’ value of 1; Fig. 3B). The parasites had for the most part a low degree, and the distribution of the specialisation index indicated that several of the parasites were specialists (*d*’ values ~1; Fig. 3D). The parasites showed the highest relative number of specialised taxa in PIDA. However, the parasites also included the taxa with the highest degree (Fig. 3C), the well-known parasites belonging to the Syndiniales (MALV II) and Perkinsidae.

### Interlinked species

Interlinked species^24^ are taxa present in either several types of interactions, or on both sides of the same interaction. An interlinked species is for example a species that is registered as a predator in the predator-prey network, and is also present as a host in the symbiont-host network. In total there were 117 interlinked species in PIDA (~5% of the total entries in PIDA, Fig. 4; Table 2). The maximum number of interaction types for any species was three (Table 2; Panel A-E). The majority of interlinked species occurred in the overlap of species recorded as predator, as prey and as host of parasites (Table 2; panel A). The interlinked species that held a role in each of the three independent bipartite networks (i.e. in the predator-prey and symbiont-host and in the parasite-host) are shown in panel B and E in Table 2 (corresponding to the overlap B and E in Fig. 4). The majority of interlinked species were represented with two interaction types (Table 2; panel F-N). There was also an example of hyperparasitism where one species had the role of parasite for another organism as well as being hosts of other parasites (Table 2; panel I).

**Fig. 4.**
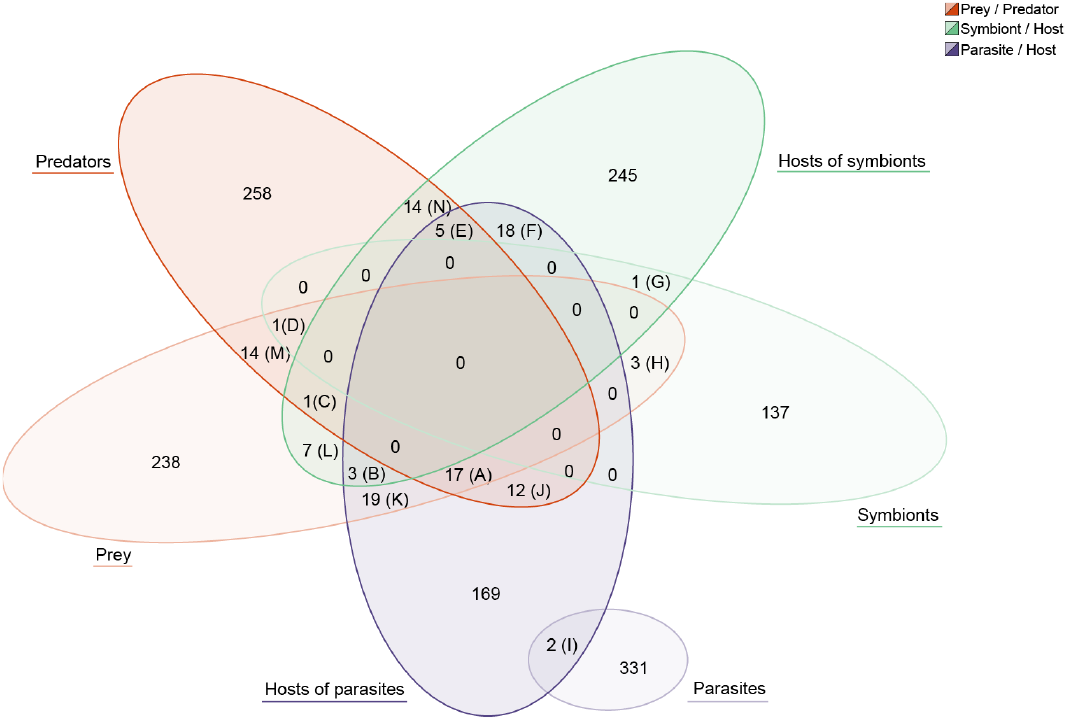
Interlinked species. Venn diagram illustrating the number of species in PIDA that hold multiple ‘roles’ and are present in more than one interaction network/type; i.e. either present in two distinct networks (e.g. registered as ‘Prey’ in the predator-prey network, and also as ‘Symbiont’ in the symbiont-host network) or on both sides of the same network (e.g. registered as both ‘Predator’ and ‘Prey’ in the predator-prey network). Letters A – N refers to the panels in Table 2 where the taxonomy of the different overlapping species and their roles are presented. ‘Parasites’ were only found to overlap with ‘Host of parasites’, representing two cases of hyperparasitism (i.e. parasite species parasitizing other parasite species). Only taxa with full species name determined were included to avoid overestimating overlapping species (e.g. *Amoebophrya* NA and similar were excluded).

**Table 2:** Interlinked species in PIDA. Panel A-N refers to the 14 overlapping sections in the Venn diagram (Fig. 4). The taxonomy of the overlapping species in PIDA is listed at species, phylum and supergroup level. The column “No” refers to the number of roles the overlapping species held, i.e. the number of bipartite networks (or interaction types in the same network) the species occurred in. The column “Present as” displays which interaction

| Panel | Species | Phylum | Supergroup | No | Present as |  |  |
| --- | --- | --- | --- | --- | --- | --- | --- |
| A | *Akashiwo sanguinea* | Dinoflagellata | Alveolata | 3 | Predator | Prey | Host of par |
| A | *Alexandrium catenella* | Dinoflagellata | Alveolata | 3 | Predator | Prey | Host of par |
| A | *Alexandrium ostenfeldii* | Dinoflagellata | Alveolata | 3 | Predator | Prey | Host of par |
| A | *Alexandrium tamarense* | Dinoflagellata | Alveolata | 3 | Predator | Prey | Host of par |
| A | *Cochlodinium polykrikoides* | Dinoflagellata | Alveolata | 3 | Predator | Prey | Host of par |
| A | *Dinophysis acuminata* | Dinoflagellata | Alveolata | 3 | Predator | Prey | Host of par |
| A | *Gymnodinium catenatum* | Dinoflagellata | Alveolata | 3 | Predator | Prey | Host of par |
| A | *Gymnodinium sanguineum* | Dinoflagellata | Alveolata | 3 | Predator | Prey | Host of par |
| A | *Heterocapsa rotundata* | Dinoflagellata | Alveolata | 3 | Predator | Prey | Host of par |
| A | *Heterocapsa triquetra* | Dinoflagellata | Alveolata | 3 | Predator | Prey | Host of par |
| A | *Neoceratium furca* | Dinoflagellata | Alveolata | 3 | Predator | Prey | Host of par |
| A | *Oblea rotunda* | Dinoflagellata | Alveolata | 3 | Predator | Prey | Host of par |
| A | *Oxyrrhis marina* | Dinoflagellata | Alveolata | 3 | Predator | Prey | Host of par |
| A | *Proocentrum micans* | Dinoflagellata | Alveolata | 3 | Predator | Prey | Host of par |
| A | *Proocentrum minimum* | Dinoflagellata | Alveolata | 3 | Predator | Prey | Host of par |
| A | *Protoperdinium pellucidum* | Dinoflagellata | Alveolata | 3 | Predator | Prey | Host of par |
| A | *Scrippsiella trochoidea* | Dinoflagellata | Alveolata | 3 | Predator | Prey | Host of par |
| B | *Guinardia delicatula* | Diatomea | Stramenopiles | 3 | Prey | Host of par | Host of symb |
| B | *Thalassiosira rotula* | Diatomea | Stramenopiles | 3 | Prey | Host of par | Host of symb |
| B | *Paramecium bursaria* | Ciliophora | Alveolata | 3 | Prey | Host of par | Host of symb |
| C | *Euplotes aediculatus* | Ciliophora | Alveolata | 3 | Predator | Prey | Host of symb |
| D | *Heterocapsa rotunda* | Dinoflagellata | Alveolata | 3 | Predator | Prey | Symbiont |
| E | *Acanthamoeba castellanii* | Discosea | Amoebozoa | 3 | Predator | Host of par | Host of symb |
| E | *Acanthamoeba polyphaga* | Discosea | Amoebozoa | 3 | Predator | Host of par | Host of symb |
| E | *Euplotes woodruffi* | Ciliophora | Alveolata | 3 | Predator | Host of par | Host of symb |
| E | *Noctiluca scintillans* | Dinoflagellata | Alveolata | 3 | Predator | Host of par | Host of symb |
| E | *Thalassicolla nucleata* | Polycystinea | Rhizaria | 3 | Predator | Host of par | Host of symb |
| F | *Saccamoeba lacustris* | Tubulinea | Amoebozoa | 2 | Host of par | Host of symb | - |
| F | *Pseudo-nitzschia pungens* | Diatomea | Stramenopiles | 2 | Host of par | Host of symb | - |
| F | *Durinskia baltica* | Dinoflagellata | Alveolata | 2 | Host of par | Host of symb | - |
| F | *Durinskia dybowskii* | Dinoflagellata | Alveolata | 2 | Host of par | Host of symb | - |
| F | *Eutintinnus tenuis* | Ciliophora | Alveolata | 2 | Host of par | Host of symb | - |
| F | *Kryptoperidinium foliaceum* | Dinoflagellata | Alveolata | 2 | Host of par | Host of symb | - |
| F | *Neoceratium horridum* | Dinoflagellata | Alveolata | 2 | Host of par | Host of symb | - |
| F | *Podolampas bipes* | Dinoflagellata | Alveolata | 2 | Host of par | Host of symb | - |
| F | *Spirostomum ambiguum* | Ciliophora | Alveolata | 2 | Host of par | Host of symb | - |
| F | *Stentor polymorphus* | Ciliophora | Alveolata | 2 | Host of par | Host of symb | - |
| F | *Phorticium pylonium* | Polycystinea | Rhizaria | 2 | Host of par | Host of symb | - |
| F | *Acanthometra pellucida* | Acantharia | Rhizaria | 2 | Host of par | Host of symb | - |
| F | *Acanthostaurus purpurascens* | Acantharia | Rhizaria | 2 | Host of par | Host of symb | - |
| F | *Collozoum caudatum* | Polycystinea | Rhizaria | 2 | Host of par | Host of symb | - |
| F | *Collozoum inerme* | Polycystinea | Rhizaria | 2 | Host of par | Host of symb | - |
| F | *Collozoum pelagicum* | Polycystinea | Rhizaria | 2 | Host of par | Host of symb | - |
| F | *Hexacantium pachydermum* | Polycystinea | Rhizaria | 2 | Host of par | Host of symb | - |
| F | *Thalassicolla spumida* | Polycystinea | Rhizaria | 2 | Host of par | Host of symb | - |
| G | *Akashiwo sanguineum* | Dinoflagellata | Alveolata | 2 | Host of symb | Symbiont | - |
| H | *Chlamydomonas hedleyi* | Chlorophyceae | Archaeplastida | 2 | Prey | Symbiont | - |
| H | *Nitzschia frustulum* | Diatomea | Stramenopiles | 2 | Prey | Symbiont | - |
| H | *Protodinium simplex* | Dinoflagellata | Alveolata | 2 | Prey | Symbiont | - |
| I | *Keppenodinium mycetoides* | Syndiniales | Alveolata | 2 | Host of par | Parasite | - |
| I | *Oodinium acanthometrae* | Dinoflagellata | Alveolata | 2 | Host of par | Parasite | - |
| J | *Dinophysis acuta* | Dinoflagellata | Alveolata | 2 | Predator | Host of par | - |
| J | *Dinophysis norvegica* | Dinoflagellata | Alveolata | 2 | Predator | Host of par | - |
| J | *Diplopsalis lenticula* | Dinoflagellata | Alveolata | 2 | Predator | Host of par | - |
| J | *Karlodinium veneficum* | Dinoflagellata | Alveolata | 2 | Predator | Host of par | - |
| J | *Gonyaulax polygramma* | Dinoflagellata | Alveolata | 2 | Predator | Host of par | - |
| J | *Protoperidinium bipes* | Dinoflagellata | Alveolata | 2 | Predator | Host of par | - |
| J | *Protoperidinium minutum* | Dinoflagellata | Alveolata | 2 | Predator | Host of par | - |
| J | *Protoperidinium steinii* | Dinoflagellata | Alveolata | 2 | Predator | Host of par | - |
| J | *Eutintinnus pectinis* | Ciliophora | Alveolata | 2 | Predator | Host of par | - |
| J | *Favella ehrenbergii* | Ciliophora | Alveolata | 2 | Predator | Host of par | - |
| J | *Trithigmostoma cucullulus* | Ciliophora | Alveolata | 2 | Predator | Host of par | - |
| J | *Strombidium capitatum* | Ciliophora | Alveolata | 2 | Predator | Host of par | - |
| K | *Coscinodiscus granii* | Diatomea | Stramenopiles | 2 | Prey | Host of par | - |
| K | *Coscinodiscus radiatus* | Diatomea | Stramenopiles | 2 | Prey | Host of par | - |
| K | *Cylindrotheca closterium* | Diatomea | Stramenopiles | 2 | Prey | Host of par | - |
| K | *Eucampia zoodiacus* | Diatomea | Stramenopiles | 2 | Prey | Host of par | - |
| K | *Guinardia flaccida* | Diatomea | Stramenopiles | 2 | Prey | Host of par | - |
| K | *Odontella sinensis* | Diatomea | Stramenopiles | 2 | Prey | Host of par | - |
| K | *Stephanopyxis turris* | Diatomea | Stramenopiles | 2 | Prey | Host of par | - |
| K | *Thalassionema nitzschioides* | Diatomea | Stramenopiles | 2 | Prey | Host of par | - |
| K | *Thalassiosira nordenskiöldii* | Diatomea | Stramenopiles | 2 | Prey | Host of par | - |
| K | *Chaetoceros didymus* | Diatomea | Stramenopiles | 2 | Prey | Host of par | - |
| K | *Leptocylinthus danicus* | Diatomea | Stramenopiles | 2 | Prey | Host of par | - |
| K | *Thalassiosira punctigera* | Diatomea | Stramenopiles | 2 | Prey | Host of par | - |
| K | *Gymnodinium instriatum* | Dinoflagellata | Alveolata | 2 | Prey | Host of par | - |
| K | *Gymnodinium mikimotoi* | Dinoflagellata | Alveolata | 2 | Prey | Host of par | - |
| K | *Gyrodinium aureolum* | Dinoflagellata | Alveolata | 2 | Prey | Host of par | - |
| K | *Neoceratium fusus* | Dinoflagellata | Alveolata | 2 | Prey | Host of par | - |
| K | *Neoceratium lineatum* | Dinoflagellata | Alveolata | 2 | Prey | Host of par | - |
| K | *Neoceratium tripos* | Dinoflagellata | Alveolata | 2 | Prey | Host of par | - |
| K | *Paramecium tetraurelia* | Ciliophora | Alveolata | 2 | Prey | Host of par | - |
| L | *Corethron hystris* | Diatomea | Stramenopiles | 2 | Prey | Host of symb | - |
| L | *Skeletonema costatum* | Diatomea | Stramenopiles | 2 | Prey | Host of symb | - |
| L | *Thalassiosira pseudonana* | Diatomea | Stramenopiles | 2 | Prey | Host of symb | - |
| L | *Euplotes octocarinatus* | Ciliophora | Alveolata | 2 | Prey | Host of symb | - |
| L | *Eutintinnus tubulosus* | Ciliophora | Alveolata | 2 | Prey | Host of symb | - |
| L | *Paramecium caudatum* | Ciliophora | Alveolata | 2 | Prey | Host of symb | - |
| L | *Volvox carteri* | Chlorophyceae | Archaeplastida | 2 | Prey | Host of symb | - |
| M | *Fibrocapsa japonica* | Raphidophyceae | Stramenopiles | 2 | Predator | Prey | - |
| M | *Heterosigma akashiwo* | Raphidophyceae | Stramenopiles | 2 | Predator | Prey | - |
| M | *Pseudobodo tremulans* | Bicosoecida | Stramenopiles | 2 | Predator | Prey | - |
| M | *Archerella flavum* | Labyrinthulomy | Stramenopiles | 2 | Predator | Prey | - |
| M | *Favella taraikaensis* | Ciliophora | Alveolata | 2 | Predator | Prey | - |
| M | *Amphidinium carterae* | Dinoflagellata | Alveolata | 2 | Predator | Prey | - |
| M | *Lingulodinium polyedrum* | Dinoflagellata | Alveolata | 2 | Predator | Prey | - |
| M | *Myrionecta rubra* | Ciliophora | Alveolata | 2 | Predator | Prey | - |
| M | *Pfiesteria piscicida* | Dinoflagellata | Alveolata | 2 | Predator | Prey | - |
| M | *Prorocentrum triestinum* | Dinoflagellata | Alveolata | 2 | Predator | Prey | - |
| M | *Bodo saltans* | Kinetoplastea | Excavata | 2 | Predator | Prey | - |
| M | *Rhyncomonas nasuta* | Kinetoplastea | Excavata | 2 | Predator | Prey | - |
| M | *Prymnesium parvum* | Prymnesiophyceae | Haptophyta | 2 | Predator | Prey | - |
| M | *Chrysochromulina polylepis* | Prymnesiophyceae | Haptophyta | 2 | Predator | Prey | - |
| N | *Spirostomum minus* | Ciliophora | Alveolata | 2 | Predator | Host of symb | - |
| N | *Strombidium sulcatum* | Ciliophora | Alveolata | 2 | Predator | Host of symb | - |
| N | *Euplotes daidaleos* | Ciliophora | Alveolata | 2 | Predator | Host of symb | - |
| N | *Climacostomum virens* | Ciliophora | Alveolata | 2 | Predator | Host of symb | - |
| N | *Peridinium quinquecome* | Dinoflagellata | Alveolata | 2 | Predator | Host of symb | - |
| N | *Levanderina fissa* | Dinoflagellata | Alveolata | 2 | Predator | Host of symb | - |
| N | *Orbulina universa* | Foraminifera | Rhizaria | 2 | Predator | Host of symb | - |
| N | *Pulleniatina obliquiloculata* | Foraminifera | Rhizaria | 2 | Predator | Host of symb | - |
| N | *Sorites marginalis* | Foraminifera | Rhizaria | 2 | Predator | Host of symb | - |
| N | *Globigerinoides ruber* | Foraminifera | Rhizaria | 2 | Predator | Host of symb | - |
| N | *Globigerinoides sacculifer* | Foraminifera | Rhizaria | 2 | Predator | Host of symb | - |
| N | *Globorotalia menardii* | Foraminifera | Rhizaria | 2 | Predator | Host of symb | - |
| N | *Archaias angulatus* | Foraminifera | Rhizaria | 2 | Predator | Host of symb | - |
| N | *Lenisia limosa* | Breviatea | Obazoa | 2 | Predator | Host of symb | - |

### The SAR supergroup: dominance in literature vs. dominance in environmental studies

The SAR supergroup heavily dominated PIDA, and we therefore examined this supergroup more in depth. Since SAR is also known to dominate environmental sequencing studies^25–27^, we compared the SAR records in PIDA with one of the most well-known recent environmental diversity studies; the *Tara* Oceans survey^25^. We identified taxonomic groups that were found in high diversity and abundance in the *Tara* Oceans survey, but that had few entries in PIDA (yellow circles in Fig. 5). Within Stramenopiles the groups that appeared to be especially underrepresented compared with environmental sequencing data were the Labyrinthulomycetes and Marine Stramenopiles (MASTs). These groups were represented by ~1,100 Operational Taxonomic Units (OTUs, a species proxy) in the environmental study by de Vargas et al.^25^, while comprising only 12 entries in PIDA (~0.5% of the total entries). The most prominent underrepresentation of the alveolates was Syndiniales (MALV II). Even though the Syndiniales were relatively numerous in PIDA (200 entries; ~8% of the total entries), the MALV II/Syndiniales comprised astonishing ~5,600 OTUs in the study by de Vargas et al.^25^ Within Rhizaria the foraminifers were fairly well represented in PIDA (330 entries, >55 unique species) compared to *Tara* Oceans (~250 OTUs) and also compared to a recent study of Morard et al^28^. The other rhizarian groups such as Acantharia, Polycystinea and Cercozoa were, however, poorly represented in PIDA compared to the diversity in the *Tara* Oceans study (~100-150 entries with ~1,000 to >5,000 OTUs in *Tara).* Apicomplexa did not comprise many entries in PIDA because these parasites only infect multicellular (metazoan) hosts. Therefore, the few apicomplexans that are present in PIDA are recorded as hosts of parasites, symbionts or as prey.

**Fig. 5.**
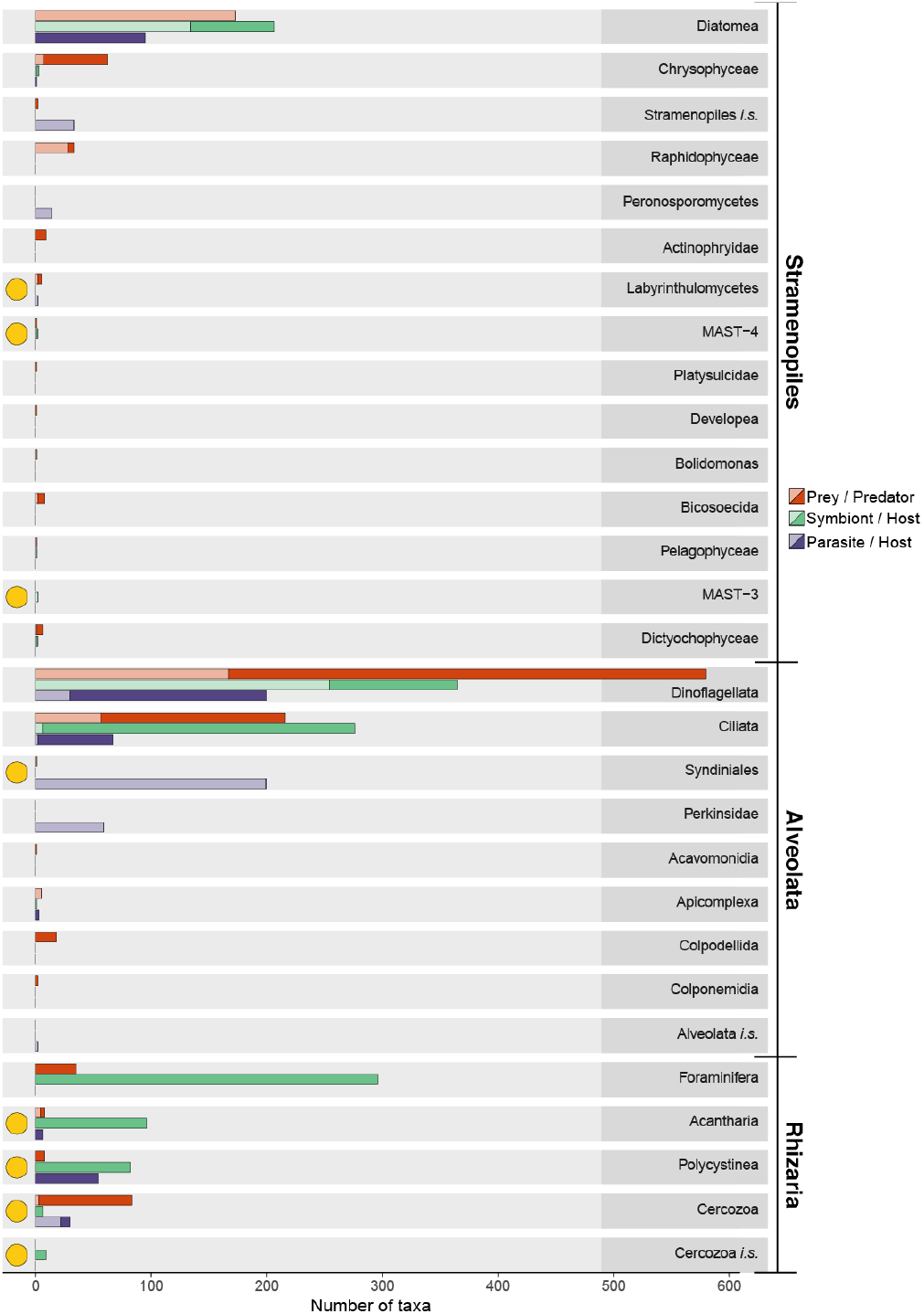
The dominating SAR supergroup. Number of interactions registered in the PIDA database belonging to the SAR supergroup (Stramenopiles, Alveolates and Rhizaria). For each of the SAR supergroups the entries of the different taxonomic groups at ‘phylum level’ are shown (corresponding to the third taxonomic level in PIDA). Red bars show predators, green represent symbiosis and purple parasitism. Solid colours represent host/predator and transparent colours represent symbiont/parasite/prey. Yellow circles highlight the ‘phyla’ that comprise few records in PIDA compared to the (hyper)diversity these ‘phyla’ represent in environmental HTS studies, such as the TARA ocean study^25^. Abbreviations used in this figure: *i.s.* refers to *Incertae sedis* or unknown.

## Discussion

Our comprehensive bibliographic survey shows that microbial interactions are spread across all major eukaryotic groups as well as across the main bacterial (e.g. Cyanobacteria, Bacteroidetes, Proteobacteria and Firmicutes) and archaeal (Halobacteria, Methanobacteria and Methanomicrobia) lineages. All major protistan groups were involved in interactions in one or multiple ways; as hosts, symbionts, parasites, predators and/or prey (Fig. 1 A-D). But our data showed that there is a bias which is especially skewed towards well-known representatives belonging to the SAR supergroup (i.e., Stramenopiles, Alveolata and Rhizaria). Members of the SAR supergroup have historically gained much attention in the protist research community since they are key species in marine ecosystems, holding important roles as primary producers, parasites, symbionts and predators/grazers^25^. Within SAR the distribution was further skewed towards certain well-characterized and species-rich lineages, such as dinoflagellates, ciliates, and diatoms. In particular, the dinoflagellate and diatom lineages include many species that hold key roles as primary producers and CO_2_ sequesters in the ocean^29^, and have consequently been the subject of many studies. Dinoflagellates also include species that can form harmful algal blooms (HABs)^30^, producing toxins that can have devastating effects on fisheries and aquaculture, making the research on their ecology and life cycles a priority^31^. Other heterotrophic dinoflagellates and ciliates are among the most important predators in aquatic ecosystems, consuming a significant proportion of both bacterial and protist biomass^32,33^. Although the SAR supergroup dominated PIDA, our comparison of the SAR records with the *Tara* Oceans data^25^ revealed that there are several SAR lineages, which are abundant and diverse in the marine realm, that were underrepresented when it comes to characterization of their ecological roles as interactors in aquatic environments. Compared to environmental studies there were also several other protist lineages that were underrepresented in PIDA. Fungi, for example, have been shown to be diverse in environmental HTS surveys of several aquatic environments^34,35^ and some of these have been proposed as important parasites of protists^36,37^. Judging from the scant number of entries in PIDA and the relatively broad host ranges that these organisms had, we suggest that future investigations should focus attention on revealing more about the ecological function of aquatic fungi. Another example of an underrepresented lineage in PIDA is Excavata. In the *Tara* Oceans surveys the enigmatic diplonemids were shown to be truly hyperdiverse, with more than 12.000 OTUs, while they have a meagre four entries in PIDA. It was not surprising that diplonemids were poorly represented in PIDA since their immense diversity was only recently discovered^25^, and because so little is yet known about the lifestyle of these excavates^38^. But it underlines that diplonemids, and other excavates represents a black box also when it comes to protist interactions.

Protist predation or grazing is crucial for channelling carbon and energy to higher trophic levels^39^ as well as for the release of dissolved nutrients to the base of the food web^40^. The bipartite network analysis of the predator-prey interactions in PIDA indicated that predation as an ecological strategy is not directed towards either specialisation nor generalisation, but that predators are “multivorous” and feed on several different prey organisms. But instead of hunting for prey many organisms depend on other strategies for resource acquisition that involve more intimate relationships, such as the interaction between parasites or symbionts and their hosts. These intimate interactions have evolved independently from free-living ancestors multiple times in diverse and evolutionary unrelated protist lineages. To develop such intimate interactions requires a high degree of specialization as it necessitates a metabolic dialogue between the interacting organisms. Our bipartite network analyses for symbiont-host and parasite-host interactions showed that many of the symbiont and parasite species seemed to be moderately specialized.

The importance and scientific relevance of symbiosis was reflected by the great variety of symbiotic interaction we found. Several protists harbour microbial symbionts (eukaryotic and prokaryotic) that provide e.g. carbohydrates (through photosynthesis), vitamins, Nitrogen (through N_2_-fixation) and defence to their hosts, in exchange for other nutrients, vitamins and protection^1,4^. An example of this is Cyanobacteria that live inside their protist hosts (e.g., diatoms, dinoflagellates or radiolarians) and provide photosynthesis products^41,42^ or nitrogen through nitrogen fixation^43–45^, in exchange for protection and/or nutrients. There were also many records of heterotrophic bacteria engaged in symbiosis with protists in PIDA, where bacteria provide their hosts with vitamins and other types of nutrients in exchange for photosynthesis products (carbohydrates), nutrients, or protection. Such types of symbiotic interactions have been demonstrated for several relationships between bacteria and microalgae^46–48^. One example is the relationship between the diatom *Pseudo-nitzschia multiseries* and *Sulfitobacter* sp. SA11. *Pseudo-nitzschia multiseries* secretes organic carbon and a sulphonated metabolite called taurine which is taken up by the *Sulfitobacter* sp., and the *Sulfitobacter* sp. bacteria respond by secreting ammonium for the diatom and then switch their preference from ammonium to nitrate, thereby promoting the growth rate of both partners involved in the symbiosis^49^. There were remarkably few studies demonstrating symbiotic relationships between two or more heterotrophic protists in the aquatic environment. The only records we found all represented the same type of symbiotic relationship between the parasite *Neoparamoeba perurans* and its kinetoplastid endosymbionts. *Neoparamoeba perurans* is a well-studied organism since it causes disease in salmon, and consequently is a threat for aquaculture^50–53^.

Parasites are present in most phylogenetic groups and hold important roles in ecosystems where they can for instance alter both the structure and dynamics of food webs^54,55^. Parasites are likely largely underrepresented in studies of microbial interactions with only 18% of the entries in PIDA. This is especially prominent in the light of recent results from environmental DNA surveys, which indicate that parasites are particularly diverse and abundant in marine as well as terrestrial ecosystems^9,25,56,57^. Cercozoan parasites were only described from diatoms, although several of their close relatives are known to parasitize a plethora of macroscopic hosts^58–61^. Likewise, parasites belonging to different stramenopile lineages such as Peronosporomycetes (oomycetes), Labyrinthulomycetes and Pirsonia were also registered to mostly infect diatoms (Fig. 1D). This probably reflects that diatoms have been the subject of more scientific studies than other protist hosts, although the true diversity of parasites infecting diatoms is likely larger than what is currently known^58^.

The network analysis showed that there were some parasite species in PIDA that were registered to infect many different host species (i.e. had a broad host ranges). But in general, the majority of parasites in PIDA were registered to infect few host species, and parasite-host interactions seemed to be slightly more dominated by specialized interactions than symbiont-host interactions. For symbionts and parasites, the observed patterns could indicate that several studies have investigated these relationships from “the parasite/symbiont point of view”, and consequently well-known taxa (e.g. the parasite Amoebophrya) has been investigated more thoroughly, and a broader host range has been characterized. In contrast, several other symbionts/parasites have been detected only associated with one host, pointing to many specialized “one-to-one” relationships in the microbial world. It could however also be speculated that this is a result of detection of a parasite or symbiont by “accident”. For instance, research conducted on a group of diatoms would from time to time observe diatoms that are infected with some parasite, or that host a specific symbiont, without looking more into the host range of these parasites or symbionts (because that was not the original focus of the study). Furthermore, the search for a parasite or symbiont’s host range has until recently been like searching for a needle in a haystack. Modern molecular tools and targeted approaches have proven useful for delineating the host range of several protist parasites and will likely lead to several new discoveries in the near future.

All in all, summarizing the data on ecological interactions involving aquatic protists and other microbes from the past ~150 years allowed us to obtain a unique overview of the known interactions. Despite the biases and knowledge gaps we identified, PIDA can be used for multiple purposes, for example: 1) To identify the functional role of a microbe using taxonomically annotated environmental DNA sequences. 2) To investigate whether ecological interaction hypotheses that derive from association networks^19^ are supported by previous studies in PIDA. 3) To obtain information about the host-range of a particular parasite, the predators of a specific prey, or the symbionts from a given host. Last but not least, our work identifies knowledge gaps that should be the focus of future research.

## Materials and Methods

The *Protist Interaction DAtabase* (PIDA) was assembled between January and November 2017 through a recursive survey of papers on microbial interactions published between 1894 and 2017. The search strategy to find the relevant literature and the template for organizing the database was performed following Lima-Mendez et al.^17^. Initially, reviews resulting from the boolean search string (plankton* AND (marin* OR ocean*)) AND (parasit* OR symbios* OR mutualis*) in Scopus (https://www.scopus.com/) and Web of Science (http://webofknowledge.com/) were examined, then the references therein were further explored. In addition, literature on protist predation on other protists and bacteria were also screened. Entries from the *AquaSymbio* database (http://aquasymbio.fr/) were used to find papers not included in the initial literature search.

PIDA documents the ecological interaction between two organisms, identified down to the species level, if possible. Interactions are identified as *parasitism*, *predation* or *symbiosis* (either mutualism or commensalism). Parasitism is used in cases where the paper at hand clearly underscores a parasitic interaction. Cases of kleptoplasty together with classical predation are contained within the group of entries termed predation. Symbiosis includes endo- and ectosymbiosis and is categorized into the different forms of symbiosis (e.g. photosymbiosis).

In addition to genus and species levels, the taxonomic classification includes three additional levels chosen pragmatically to make the database more user-friendly and portable. The highest level distinguishes between eukaryotes and prokaryotes. The second level places each taxon within super groups or other high taxonomic ranks (e.g. Rhizaria or Alveolata) following the scheme of Adl et al.^21,22^. The third level places each taxon in larger groups below the supergroup taxonomic rank (mostly Phylum level; e.g. Ciliophora, Dinoflagellata and Acantharia, or Class level; e.g. Chlorophyceae, Kinetoplastea and Diplomonadida). The taxonomic names at the third level follows the nomenclature of the Silva database (release 128, May/June 2017)^62–64^ Species names have been updated to the most recent agreed-upon classification in PIDA and can therefore deviate from the original papers they stem from due to several cases of synonymization. PIDA also documents the methods used to determine the interacting species. Symbionts and/or hosts determined by any form of microscopy or direct observation are denoted 1. Symbionts and/or hosts determined by sequencing or Fluorescence In Situ Hybridization (FISH) are denoted 2. The combination of the former two is denoted 3. Most interactions with observation type 2 also have accession numbers from GenBank^65^ that are included in PIDA. A published paper is associated to each interaction entry. When a DOI is available, it is included in the database. Only interactions from aquatic systems are included (marine, brackish and freshwater). The resulting PIDA contains 2,422 entries from 528 publications and is publicly available at github (https://github.com/ramalok/PIDA).

### Bipartite networks

Bipartite networks are the representation of interactions between two distinct classes of nodes, such as plant-pollinator, parasite-host or prey-predator. Identifying structures in bipartite networks is useful in explaining their formation, function and behaviour. Here we investigated how symbiosis, parasitism and predation differ in terms of specialization (e.g., if parasite taxa have a broader host range compared to the host range of symbionts, this indicates that parasites are less specialized (and consequently more generalists) than symbionts are). All analyses were conducted in the statistical environment R v. 3.5.0. We constructed bipartite qualitative (binary) directional networks using the R-package *bipartite* v. 2.08^66^. All taxa where the taxonomy assigned to one of the “interactors” in PIDA was ‘unknown eukaryote’, ‘unidentified bacteria’ or ‘unidentified prokaryote’ were removed before further analyses of the bipartite networks. Bipartite network indices were found using the functions *networklevel*^67^ and *specieslevel*^23^ (default settings except weighted = FALSE) in the R-package *bipartite.* Bipartite networks and network analyses were performed for four taxonomic levels (‘supergroup’, ‘phylum’, genus and species). The patterns found were consistent across the taxonomic levels, therefore only the species level is shown here. Degree (number of links/edges/interactions) was calculated for prey, predators, parasites, symbionts and hosts on the species level. The specialization index *d*’ (Kullback-Leibler distance)^68^, measures the degree of specialization at the species level, and was calculated as deviation of the actual interaction frequencies from a null model that assumes all partners in the other level of the bipartite network are used in proportion to their availability. The specialization index *d*’ ranges from 0 for the most generalized to 1 for the most specialized, and was calculated for predators, host of parasites and host of symbionts.

All bar plots and density plots were constructed using the R-package *ggplot2* v. 3.1.0^69^, and the networks in Fig. 1 were visualized in CytoScape v. 3.6.1^70^.

### Interlinked species

Interlinked species were determined using the R-package *systemPipeR* v. 3.8^73^. Only taxa with full species-names determined were included to avoid overestimating overlapping species (e.g. *Amoebophrya* sp. and similar were excluded). Venn intersects were computed using the function *overLapper* and plotted using the function *vennPlot*. Parasites only overlapped with parasite hosts and were subsequently added to the Venn plot.

## Acknowledgments

This work has been supported by the projects MicroEcoSystems (Research Council of Norway 240904) as well as INTERACTOMICS (CTM2015-69936-P, MINECO, Spain) to RL. RL was supported by a Ramón y Cajal fellowship (RYC-2013-12554, MINECO, Spain). We thank Javier del Campo and Rannveig M. Jacobsen for useful suggestions and comments that helped to improve this work.

## Supplementary material

**Supplementary figure 1:**
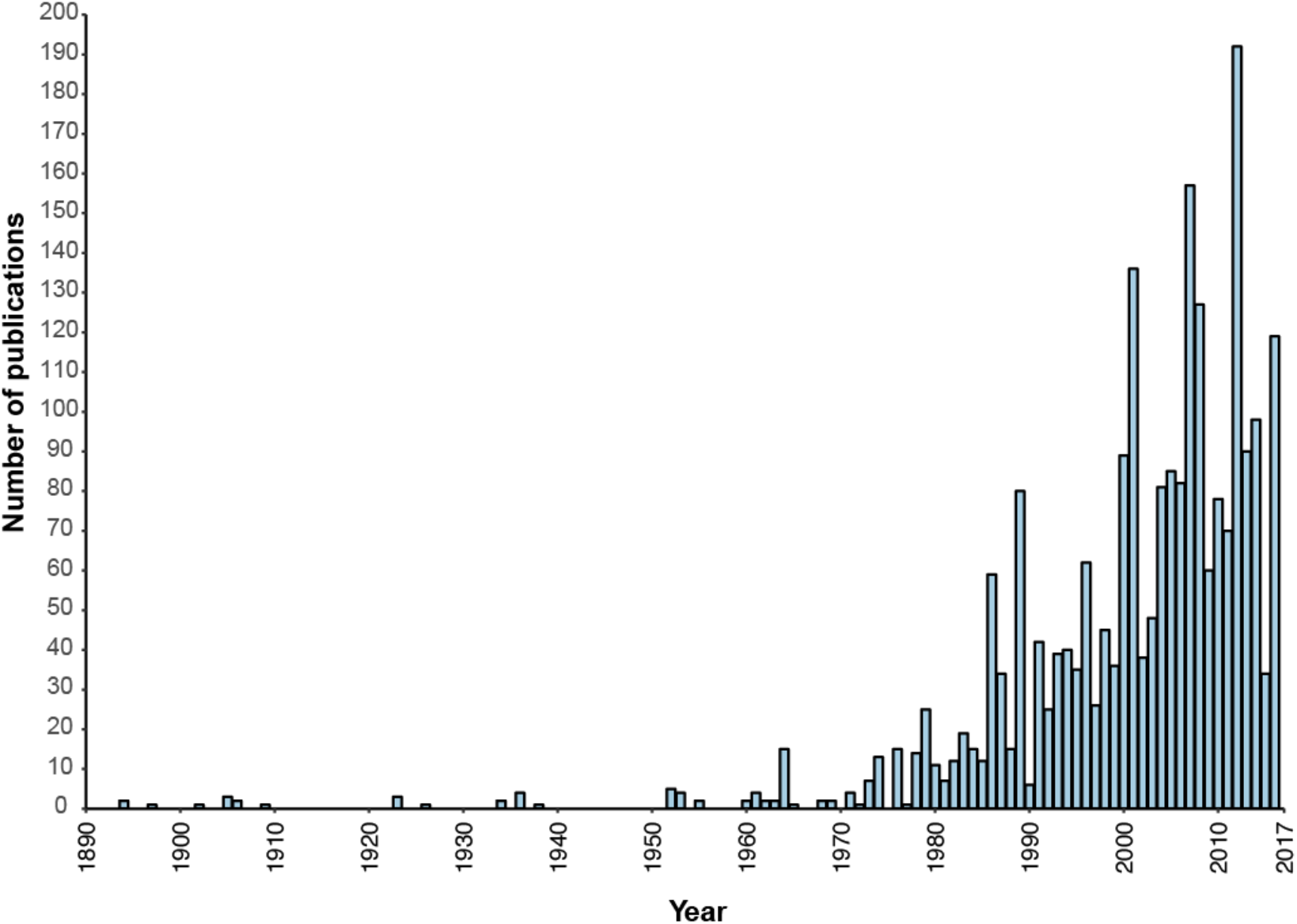
Number of scientific publications per year included in the PIDA database.

**Supplementary figure 2.**
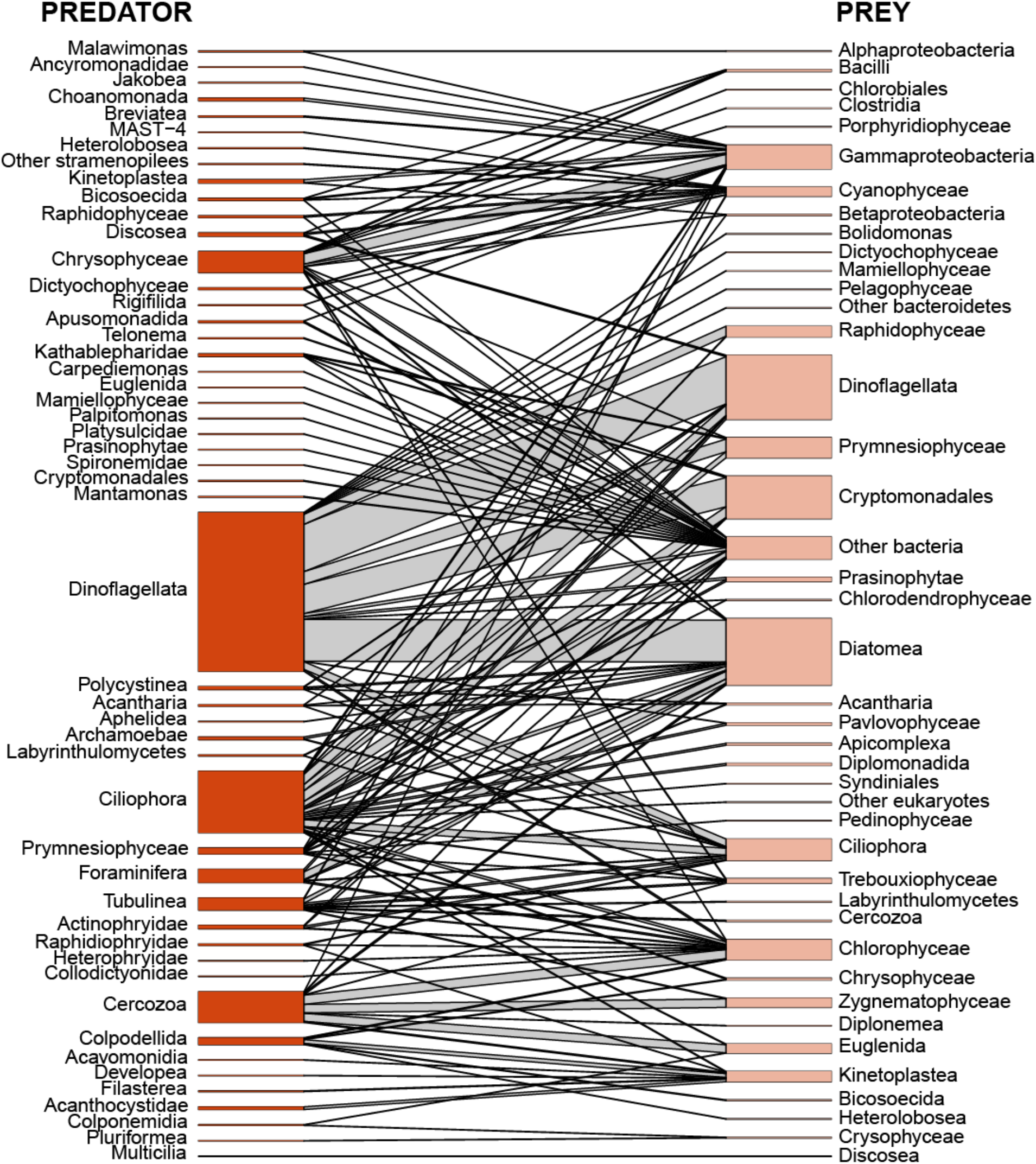
Bipartite network of Predator-Prey interactions in PIDA, at the ‘phylum level’ (corresponding to the third taxonomic level in PIDA). Predator organisms are to the left (dark orange) and prey organisms are to the right (light orange). Sizes of nodes and number of edges (i.e. lines) represent number of interactions between prey and predator taxa.

**Supplementary figure 3.**
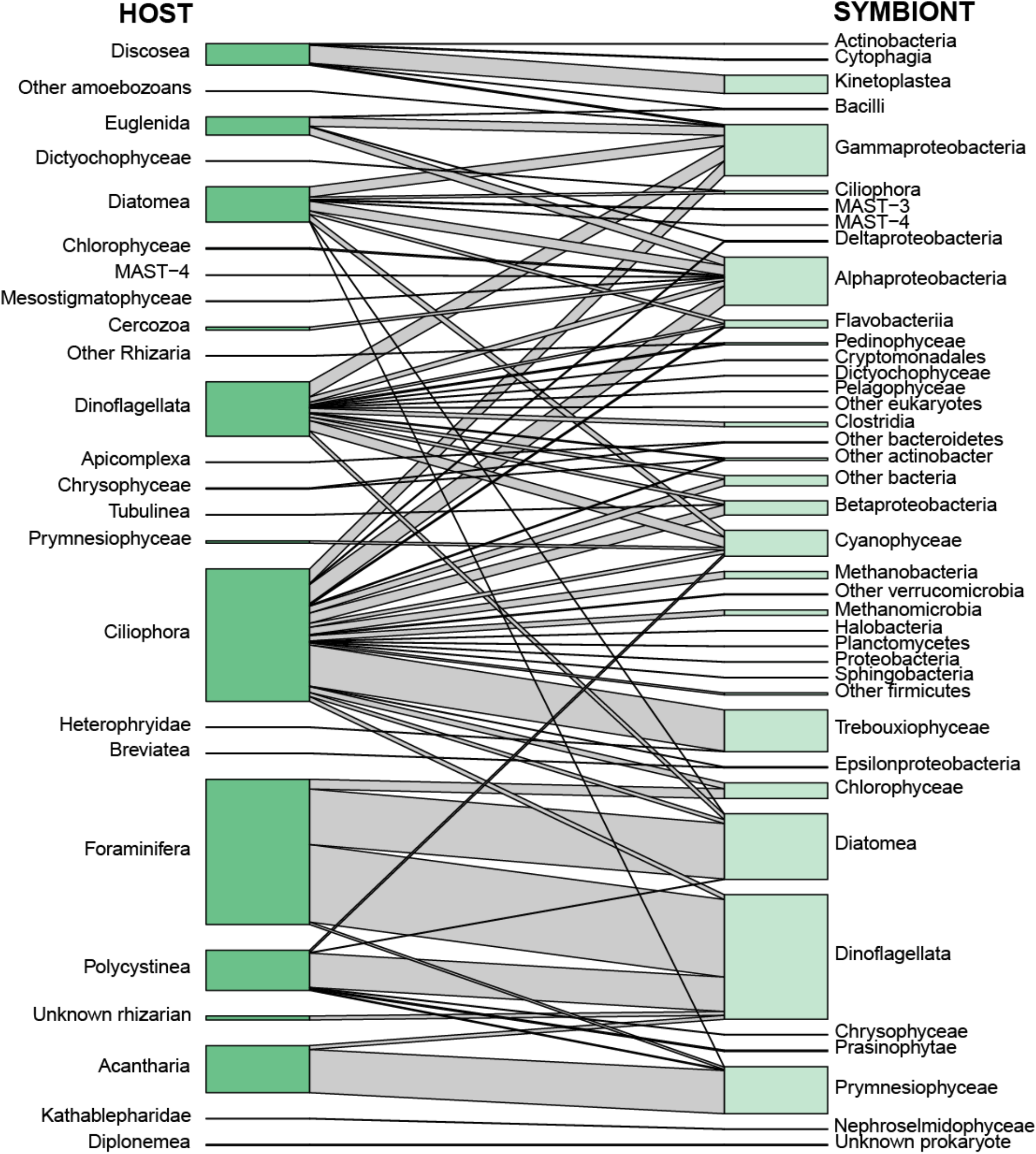
Bipartite network of Symbiont-Host interactions in PIDA, at the ‘phylum level’ (corresponding to the third taxonomic level in PIDA). Host organisms are to the left (dark green) and symbiont organisms are to the right (light green). Sizes of nodes and number of edges (i.e. lines) represent number of interactions between symbiont and host taxa.

**Supplementary figure 4.**
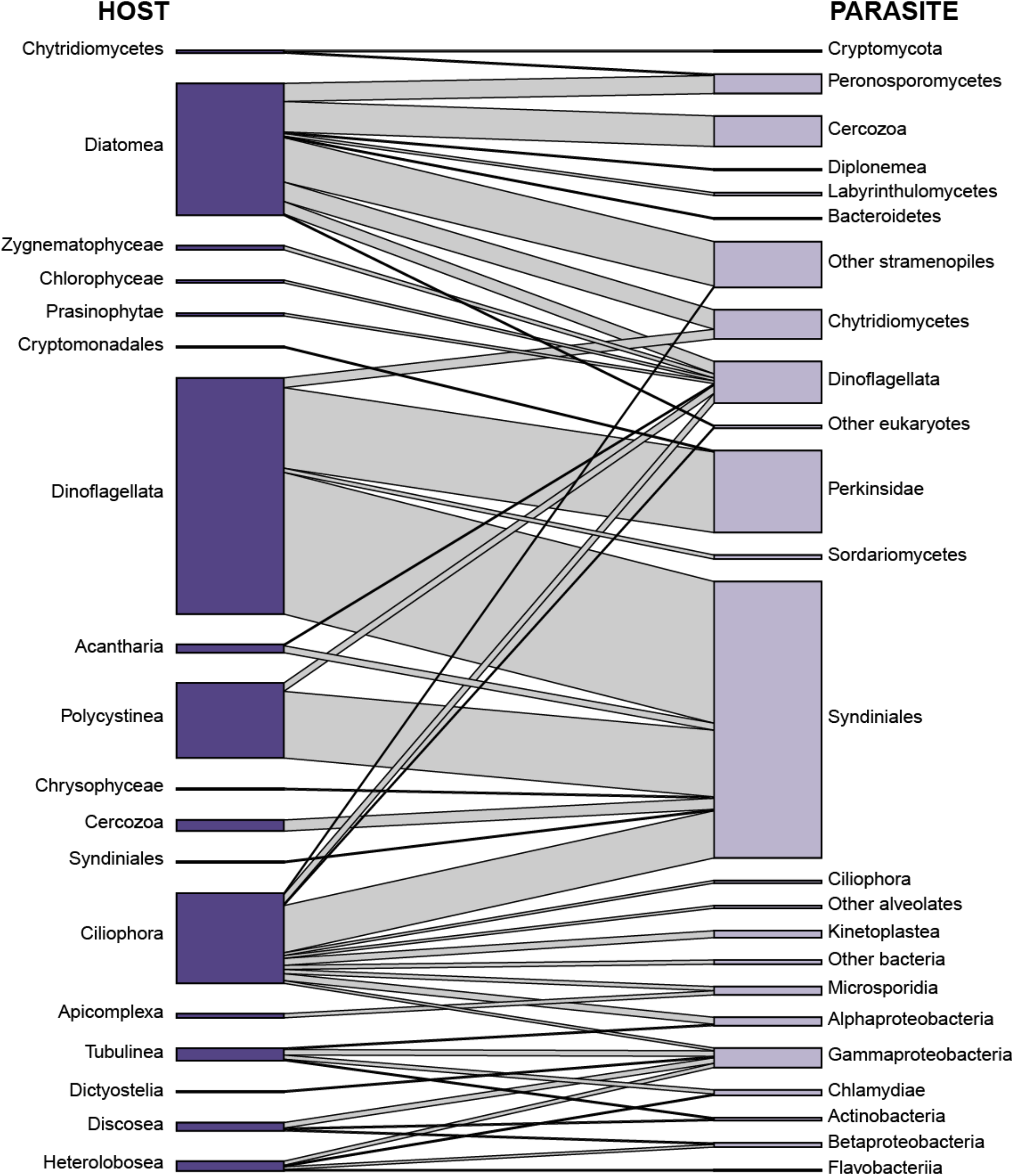
Bipartite network of Parasite-Host interactions in PIDA, at the ‘phylum level’ (corresponding to the third taxonomic level in PIDA). Host organisms are to the left (dark purple) and parasite organisms are to the right (light purple). Sizes of nodes and number of edges (i.e. lines) represent number of interactions between parasite and host taxa.

**Supplementary figure 5:**
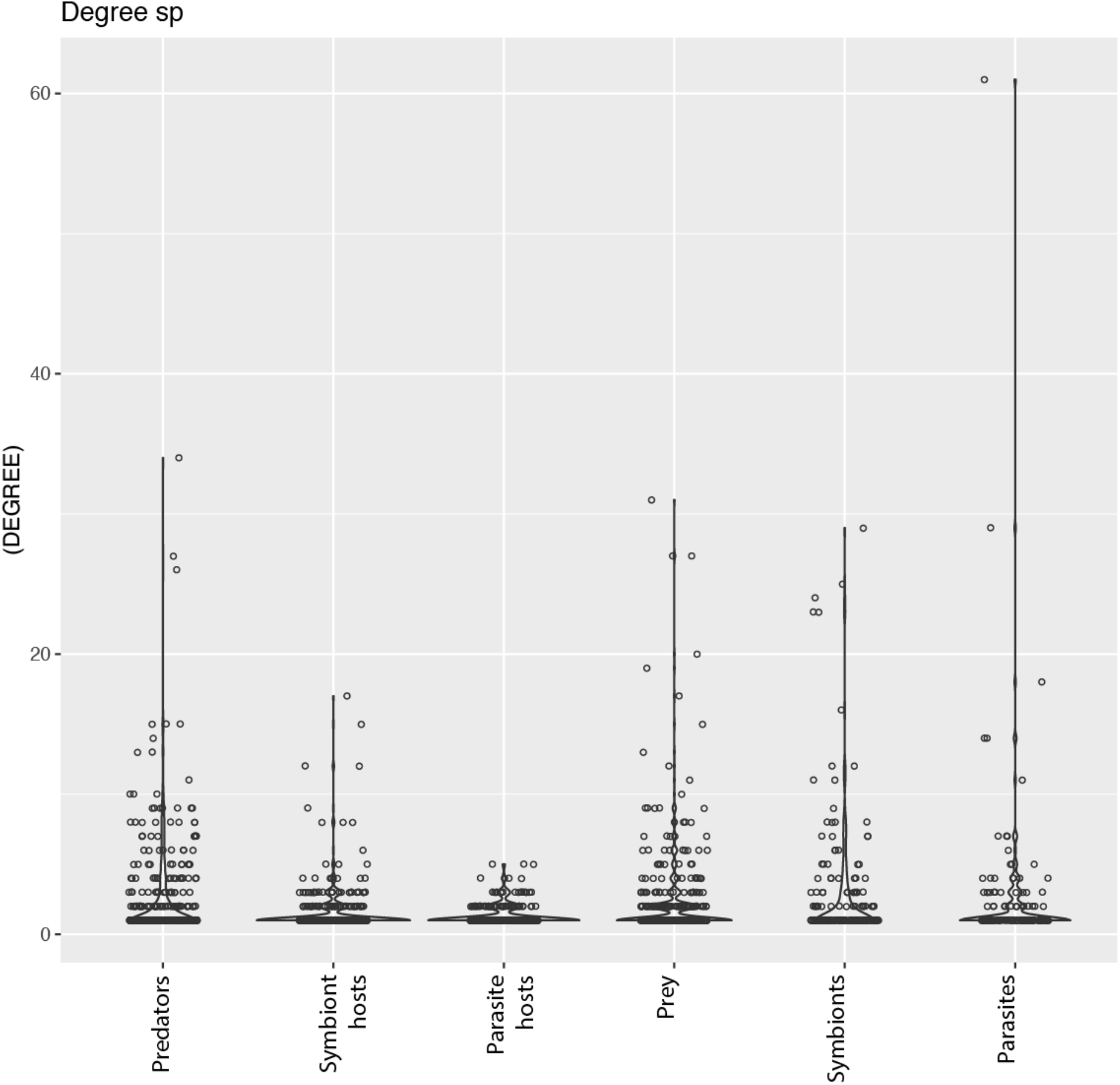
Degree displaying the number of links/edges that each species has with partners in the other level of the bipartite networks, e.g. the parasites to the far right (light purple colour) are represented with some species with high degree, i.e. registered to parasitize many different host species.

## Notes

#### Summary of Updates

-Title and abstract

https://github.com/ramalok/PIDA

